# CRISPRoff epigenetic editing for programmable gene silencing in human cells without DNA breaks

**DOI:** 10.1101/2024.09.09.612111

**Authors:** Rithu K. Pattali, Izaiah J. Ornelas, Carolyn D. Nguyen, Da Xu, Nikita S. Divekar, James K. Nuñez

## Abstract

The advent of CRISPR-based technologies has enabled the rapid advancement of programmable gene manipulation in cells, tissues, and whole organisms. An emerging platform for targeted gene perturbation is epigenetic editing, the direct editing of chemical modifications on DNA and histones that ultimately results in repression or activation of the targeted gene. In contrast to CRISPR nucleases, epigenetic editors modulate gene expression without inducing DNA breaks or altering the genomic sequence of host cells. Recently, we developed the CRISPRoff epigenetic editing technology that simultaneously establishes DNA methylation and repressive histone modifications at targeted gene promoters. Transient expression of CRISPRoff and the accompanying single guide RNAs in mammalian cells results in transcriptional repression of targeted genes that is memorized heritably by cells through cell division and differentiation. Here, we describe our protocol for the delivery of CRISPRoff through plasmid DNA transfection, as well as the delivery of CRISPRoff mRNA, into transformed human cell lines and primary immune cells. We also provide guidance on evaluating target gene silencing and highlight key considerations when utilizing CRISPRoff for gene perturbations. Our protocols are broadly applicable to other CRISPR-based epigenetic editing technologies, as programmable genome manipulation tools continue to evolve rapidly.

## 1. INTRODUCTION

CRISPR-based genome editing has revolutionized the ability to manipulate the genetic code of cells and organisms. While traditional CRISPR-Cas systems rely on inducing double-strand breaks (DSBs) or changing the sequence of the host genome, these genome editing technologies can be associated with unpredictable DNA repair outcomes, chromosomal aberrations, potential off-target editing, and cell toxicity (Doudna et al., 2020). An emerging technology to perturb gene function is to edit the epigenetic chemistry (i.e., DNA methylation and histone modifications) of the host genome that results in the modulation of gene expression without altering the underlying DNA sequence of the target gene(s). CRISPR-based epigenetic editors rely on catalytically dead Cas9 (dCas9) and the recruitment of chromatin modifiers to reprogram epigenetic marks at the targeted gene promoter that ultimately results in transcriptional repression or activation (Fig. 1).

**Figure 1.**
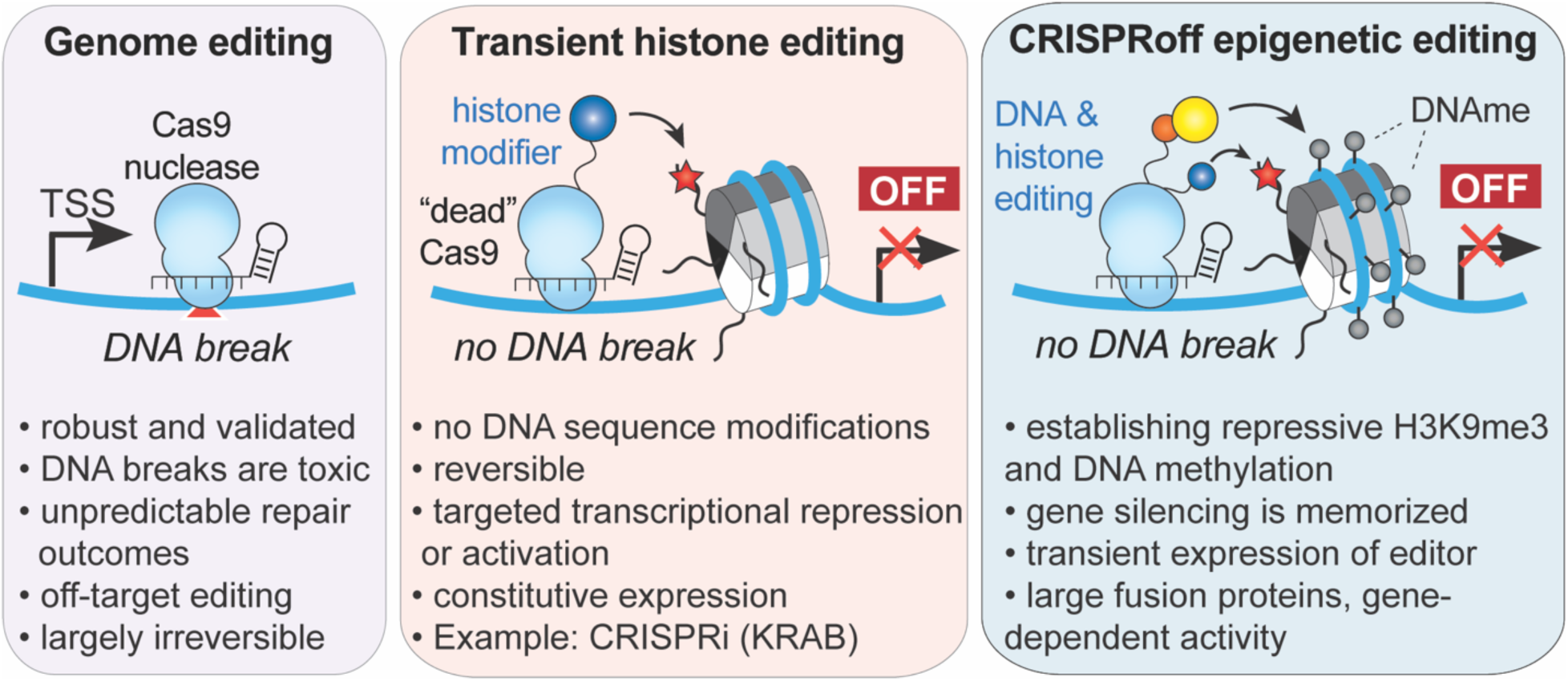
A comparison of CRISPR-based technologies for gene inactivation. While Cas9 nuclease induces double-stranded DNA breaks, epigenetic editing relies on the recruitment of repressive epigenetic modifiers to catalytically dead dCas9 to transcriptionally repress the target gene. Transient histone editing requires constitutive expression of the dCas9-epigenetic repressor to sustain gene silencing, as exemplified by CRISPRi (KRAB). CRISPRoff combinatorially programs DNA methylation and H3K9me3 at the targeted gene promoter, resulting in gene repression that is memorized and heritable across cell division, without the need for constitutive expression of CRISPRoff to sustain gene silencing.

Many CRISPR epigenetic editing technologies have been reported to date, each defined by a chromatin modifier fused to dCas9 (McCutcheon et al., 2024; Villieger et al., 2024). A commonly used epigenetic editor for transcriptional silencing is CRISPR interference (CRISPRi), which is composed of dCas9 fused to a Krüppel associated box domain (KRAB) repressor domain (Gilbert et al., 2013). KRAB serves as a scaffold for chromatin modifying proteins that establishes the histone modification H3K9me3 at the target gene promoter, leading to transcriptional repression of the target gene. For sustained target gene silencing, CRISPRi-mediated repression requires constitutive expression of dCas9-KRAB and the associated single-guide RNA (sgRNA), which is usually achieved by creating a stable cell line for *in vitro* experiments. This requirement limits its application in primary cells, complex tissues, and organisms that are challenging to engineer.

To overcome these limitations, we have developed CRISPRoff as an epigenetic editor that can heritably silence endogenous mammalian genes when targeted to their promoters without requiring constitutive expression of the CRISPR reagents (Nuñez et al., 2021). CRISPRoff is a protein fusion of dCas9 to KRAB and the DNA methyltransferase Dnmt3a-Dnmt3l (D3A-D3L) enzyme complex that establishes repressive histone H3K9me3 modifications and DNA methylation respectively at target loci for heritable transcriptional repression. Importantly, the targeted gene silencing is maintained through cell divisions after only transient expression of CRISPRoff, significantly expanding the potential for epigenetic editing treatments. Recently, the combinatorial epigenetic editing cocktail of DNMTs and KRAB have been utilized for durable gene silencing *in vitro* and *in vivo* to silence *PCSK9* and prion genes in mice (Amabile et al., 2016; Cappelluti et al., 2024; Neumann et al., 2024).

In this paper, we provide a comprehensive guide to utilizing CRISPRoff for targeted heritable gene silencing using epigenetic editing. We describe the steps involved in transiently delivering CRISPRoff into transformed cell lines, donor-isolated primary human T cells, and assessing its direct effects on transcriptional silencing. Additionally, we address potential challenges and troubleshooting strategies to ensure the successful application of this tool in various experimental contexts.

## 2. KEY CONSIDERATIONS PRIOR TO USE OF CRISPROFF

Prior to utilizing CRISPRoff for gene silencing, we recommend considering several factors:

### 2.1. Comparing epigenetic editing for gene silencing to gene knockout strategies

When deciding between CRISPRoff-mediated gene silencing and CRISPR-Cas9-mediated gene knockout or other genomic manipulation technologies such base editing and prime editing (Cong et al., 2013; Mali et al., 2013; Komor et al., 2016; Anzalone et al., 2019), it is important to consider the goals of the experiment. CRISPR-Cas9 nuclease accompanied by a sgRNA introduces genomic alterations, such as indels, at the target site to disrupt the reading frame of the gene, rendering the gene dysfunctional. Cas9 can also introduce specific mutations when combined with a donor sequence, making it ideal for experiments that require precise genome edits. However, recent studies have elucidated that CRISPR nuclease mediated knockouts can lead to genomic instabilities such as chromosomal translocations and chromothripsis, highlighting the risks of using it in mammalian cells and organisms (Adikusuma et al., 2018; Cullot et al., 2019; Fiumara et al., 2023; Kosicki et al., 2018; Tsuchida et al., 2023).

In contrast, CRISPRoff is a genetically safer alternative when the goal is to heritably silence a gene without introducing double-stranded DNA breaks, which is important when working with cells that are more susceptible to DNA damage, such as stem cells (Ihry et al., 2018) or primary cells (hematopoietic stem cells (HSCs) (Dorset et al., 2023; Fiumara et al., 2023; Lee et al., 2021) or T cells (Nahmad et al., 2023; Tsuchida et al., 2023). Lastly, epigenetic editing allows the user to reverse the programmed epigenetic changes (Morita et al., 2016). For example, CRISPRoff mediated gene repression can be reversed by actively removing DNA methylation using the TET1 demethylase enzyme, resulting in the transcriptional re-activation of the gene (Choudhury et al., 2016). This flexibility makes CRISPRoff a potent, reversible, and programmable tool for long-term gene silencing without inducing genomic alterations.

### 2.2. Delivery method of CRISPRoff

Since transient expression of CRISPRoff leads to heritable gene silencing, it circumvents the need to generate a stable cell line for constitutively expressing the CRISPRoff fusion protein. In our experience, transfection of CRISPRoff plasmids into immortalized cell lines such as HEK 293T cells and hTERT RPE-1 cells leads to durable long-term gene silencing of target genes. However, as with any DNA transfection method, we observe cellular toxicity when delivering CRISPRoff plasmid DNA to primary cells and other cell types that are sensitive to plasmid DNA. To address this, we provide an alternative protocol for synthesizing and delivering CRISPRoff *in vitro* transcribed (IVT) mRNA into specific sensitive cell types such as primary human T cells, with potential applicability for *in vivo* experiments.

### 2.3. Gene targets and cell types

While CRISPRoff can silence the majority of genes within the human genome, not all genes are amenable to CRISPRoff mediated gene silencing, owing to significant variability in the local chromatin states across different genes and different cell types. For example, CRISPRoff requires CpG dinucleotides for DNA methylation, often found as CpG islands of gene promoters (Deaton et al., 2011). Previously, we discovered that genes lacking annotated CpG islands can still be silenced with CRISPRoff, likely due to the presence of low levels of CpG dinucleotides that do not meet the CpG density cutoff of CpG islands (Gardiner-Garden et al., 1987). However, some genes are almost completely devoid of CpG dinucleotides and thus, are not amenable for CRISPRoff-mediated silencing (Nuñez et al., 2021). We recommend checking available genomic annotations of CpG content of the desired gene(s) to be repressed, such as by utilizing the CpG Island track in the UCSC Genome Browser, nested under the Expression and Regulation section. Additionally, the efficiency of CRISPRoff mediated gene silencing can vary between cell types. Immortalized or transformed cell lines, with their altered epigenetic landscapes, are more amenable to CRISPRoff treatment than primary cells, which may require additional optimization of editing and delivery conditions to achieve effective gene silencing.

### 2.4. Assessing efficiency and specificity of CRISPRoff

Finally, it is important to choose a quantitative method for assessing the efficiency of gene silencing after CRISPRoff treatment. We recommend quantitative PCR (qPCR), Western Blot, and/or flow cytometry to measure gene expression profiles. To assess the specificity of CRISPRoff and validate epigenetic changes, we recommend using various genomic techniques, such as RNA sequencing (RNA-seq), chromatin immunoprecipitation sequencing (ChIP-seq), and bisulfite sequencing to compare experimental and control samples.

## 3. SELECTION OF SINGLE GUIDE RNA AND CLONING

### 3.1. Design and selection of sgRNAs for CRISPRoff

The selection and design of a sgRNA is critical for ensuring the specificity and efficiency of CRISPRoff mediated gene silencing. Our previous experimental data show that active CRISPRoff sgRNAs are distributed broadly across the transcriptional start site (TSS), notably within a 1-kb window centered on the TSS of a target gene. Additionally, several bioinformatic tools (Doench et al., 2016; Horlbeck et al., 2016; Sanson et al., 2018) can be used to identify potential sgRNA sequences that minimize off-target effects while maximizing on-target activity. These tools predict sgRNA efficiency and specificity by evaluating factors such as GC content and accessibility of the target DNA as well as potential off-target sites. We recommend experimentally testing 3 to 5 different sgRNAs to determine the most optimal sgRNA sequence for any given gene. Additionally, simultaneously pooling the top 3 to 5 predicted sgRNAs targeting a gene promoter can improve the efficiency of target gene silencing.

### 3.2. Cloning sgRNA plasmid

sgRNA sequences are cloned into an expression plasmid from Addgene (ID#217306) that contains a U6 pol III promoter. The expression vector is a lentiviral plasmid that can be utilized either for constitutive sgRNA expression in the target cell line or for transient transfection alongside the CRISPRoff-expressing plasmid. In brief, the sgRNA plasmids are constructed by restriction cloning of protospacers downstream of a U6 promoter using BstXI and BlpI cut sites, as described previously (Gilbert et al., 2014). The sgRNA expression plasmids also express a T2A-mCherry reporter to allow for the assessment of sgRNA plasmid transfection efficiency. Listed below are example positive control sgRNAs that we have validated for silencing non-essential endogenous genes, along with a non-targeting negative control sgRNA.

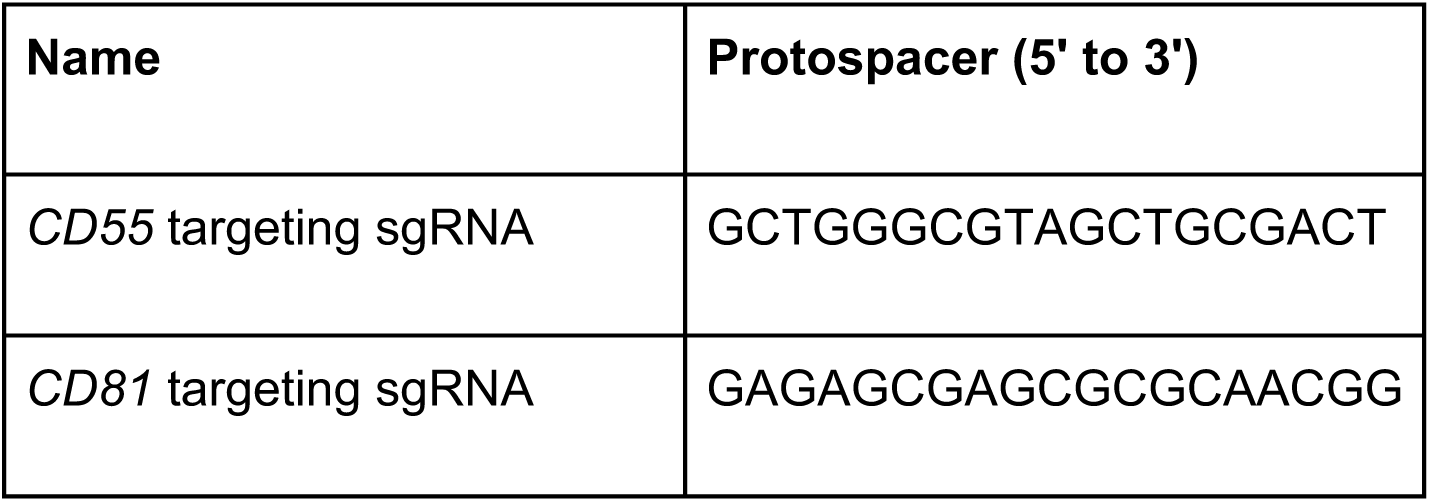
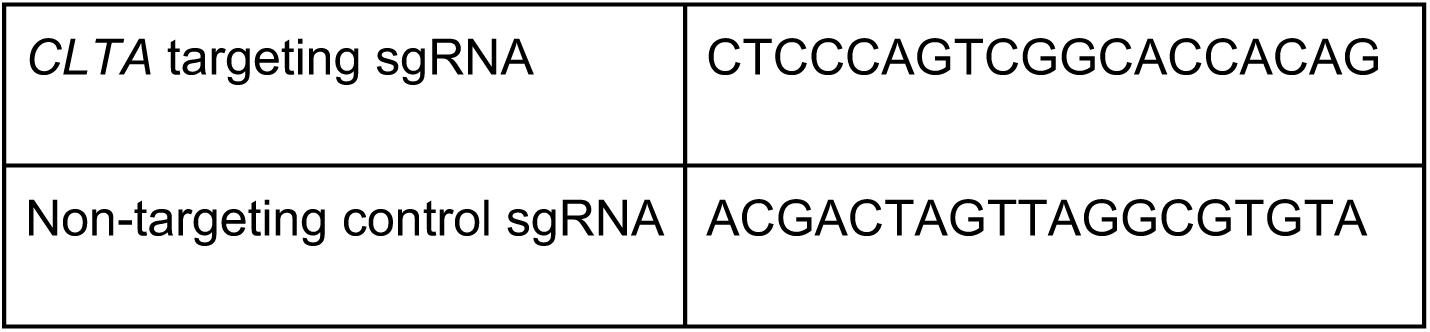

## 4. TARGETED GENE SILENCING IN HEK 293T CELLS USING CRISPROFF PLASMID

Here, we outline the process of transfecting CRISPRoff-encoding plasmid DNA into HEK 293T cells to achieve gene silencing. In this example, CRISPRoff is directed to the promoter of a non-essential gene *CLTA*, which is tagged at its endogenous genomic locus with a GFP reporter in HEK 293T cells (Leonetti et al., 2016). First, we provide a protocol for co-transfecting two plasmid constructs: one expressing CRISPRoff (fused with BFP marker) and the other expressing an sgRNA (vector encoding an mCherry marker) targeting the *CLTA* promoter. Second, we describe an alternative approach where the HEK 293T cell line constitutively expresses an sgRNA targeting the *CLTA* promoter. In this approach, only the CRISPRoff-expressing plasmid is transfected, leading to robust *CLTA* silencing. For both experiments, the readout is the silencing of CLTA-GFP that can be quantitatively measured at the protein level in single cells using flow cytometry (Fig. 2a). Other detection methods such as qPCR or Western Blot can be used for verifying the extent of CLTA knockdown at the transcript and protein levels respectively.

**Figure 2.**
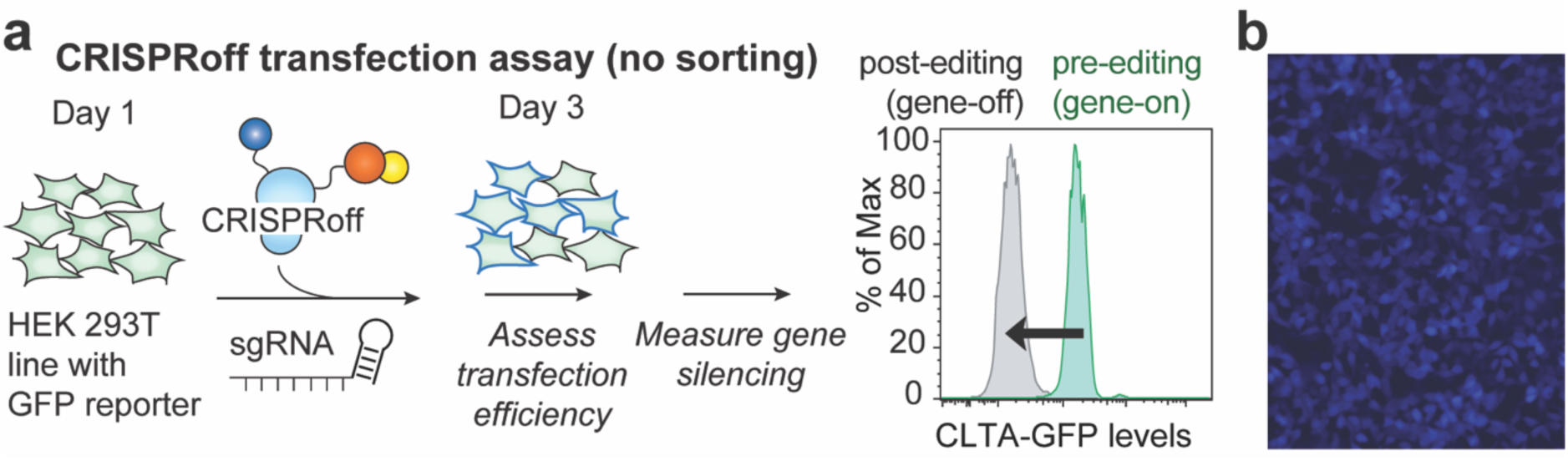
CRISPRoff plasmid transfection assay in HEK 293T cells without sorting for transfected cells. (a) The workflow for targeting CRISPRoff to the endogenous gene CLTA tagged with GFP in HEK 293T cells. Cells are first transfected with plasmid DNA-expressing CRISPRoff, measured for transfection efficiency two days post-transfection, followed by flow cytometry analysis for CLTA-GFP expression at subsequent days. (b) An example of HEK 293T cells imaged two days after transfection of CRISPRoff plasmid DNA. CRISPRoff is fused directly to a BFP fluorescent marker that can be used for quantitative measurement of transfection efficiency.

### 4.1. Required materials

- CRISPRoff expression plasmid (Addgene #167981)
- HEK 293T cells (ATCC #CRL-3216)
- Dulbecco’s modified eagle medium (DMEM) High Glucose (Thermo Fisher Scientific #11965118)
- 10% FBS (VWR #89510-186)
- Penicillin-Streptomycin-Glutamine (100X) (Gibco #1037801)
- *Trans*IT-LT1 Transfection Reagent (Mirus Bio #MIR 2225)
- Opti-MEM™ (Thermo Fisher Scientific #31985070)
- Hemocytometer or automated cell counter
- BD FACSymphony A1 Cell Analyzer (BD Biosciences)
- BD FACSAria™ Fusion Flow Cytometer (BD Biosciences)
- 24-well plates
- CO_2_ incubator
- Centrifuge
- Sterile Eppendorf tubes
- 15 ml conical tubes
- Pipettes and tips

### 4.2. Delivery of CRISPRoff by plasmid transfection

This transfection protocol has been optimized in a 24-well format. Prior to transfection, maintain HEK 293T cells in DMEM with 10% FBS and 1X Penicillin-Streptomycin-Glutamine. Passage the cells every 2–3 days, ensuring they remain at a confluency of 60-70%.

#### 4.2.1 Seeding HEK 293T cells for CRISPRoff plasmid transfection

1. Day 0: Seed HEK 293T cells to achieve ∼60 – 70% confluency the following day. We recommend seeding ∼0.9 × 10^5^ – 1.0 × 10^5^ HEK 293T cells per well in a 24-well plate in a final volume of 600 μl per well. Adjusting the seeding density might be necessary depending on the timing of transfection and the method used for cell counting. For example, seed ∼1.0 × 10^5^ – 1.1 × 10^5^ cells in the evening for transfection in the next morning; ∼0.8 × 10^5^ – 0.9 × 10^5^ cells if seeding 20-24 h before transfection.

#### 4.2.2 Transfection of CRISPRoff plasmids into HEK 293T cells

1. Day 1: Examine the cultured cells under a microscope to assess whether the cell density is approximately at a confluence of 60 – 70%.
2. Pre-warm Opti-MEM^TM^ and *Trans*IT-LT1 Transfection Reagent to room temperature for approximately 30-60 min before transfection.
3. If performing a co-transfection of CRISPRoff plasmid and an sgRNA vector, we recommend co-transfecting 300 ng of CRISPRoff plasmid and 150 ng of the target sgRNA vector into cells. CRISPRoff plasmids are diluted to be 300 ng/μl and sgRNA encoding plasmids are diluted to 150 ng/μl such that, we add 1 μl of CRISPRoff plasmid and 1 μl sgRNA vector (total of 2 μl) with the transfection reagents on Day 1.
4. Aliquot plasmid DNA into a PCR strip tube by adding:

a. 1 μl (300 ng) of CRISPRoff encoding plasmid at 300 ng/μl.
b. doing co-transfection) 1 μl (150 ng) of sgRNA vector at 150 ng/μl into the same PCR strip tube.
5. Mix 1.5 μl of the *Trans*IT-LT1 Transfection Reagent to 50 μl of pre-warmed Opti-MEM™. Incubate for 15 min at room temperature.
6. Add 51.5 μl of the *Trans*IT-LT1 Transfection Reagent-Opti-MEM™ mixture into the DNA. Incubate for 15 min at room temperature.
7. Add the mixture dropwise (∼53 μl) to cells.
8. Gently tilt the plate to evenly distribute the transfection mixture without dislodging the cells.

#### 4.2.3 Monitoring transfection using microscopy

1. Day 2: Use a fluorescent microscope to assess the transfection efficiency (Fig. 2b).

a. BFP signal (CRISPRoff) will be low or undetectable on Day 2 and will increase substantially by Day 3.
b. mCherry (sgRNA vector) signal will be high on Day 2.
2. Do not split cells or change media as this can dislodge the cells.

#### 4.2.4 Assess transfection efficiency and gene silencing by flow cytometry

1. Day 3 and beyond: Dissociate the transfected HEK 293T cells following the manufacturer’s protocol and seed cells into a fresh plate. For example, a 1:10 split into a new plate will be confluent in 2 days.
2. With dissociated cells, measure transfection efficiency (BFP and mCherry) as well as gene silencing (GFP) using the methods described in section 2.4.

a. We recommend assessing the transfection efficiency and gene silencing of CRISPRoff using flow cytometry. At Day 3, we typically detect >40% BFP+ (Fig. 3a) and >60% mCherry+ (Fig. 3b) cell populations with flow cytometry. Gene knockdown with CRISPRoff can be observed as early as Day 3, typically peaking between days 6 and 8, and in most cases, is sustained for an extended period. When collecting data over a time course (e.g., Days 3, 5, 7, 10, 14 post-transfection), we recommend normalizing the percentage of GFP-negative cells at each time point to the transfection efficiency measured on Day 3. For example, if only 50% of cells are transfected with CRISPRoff, the relative maximum percentage of cells in the population that repress the target gene is 50%. Furthermore, we recommend additional assessments of target gene silencing 2-3 weeks after transfection of CRISPRoff plasmid (Fig. 3c) to assess the durability of gene silencing.
3. If necessary, cells can be sorted for CRISPRoff and gRNA expressing cells if a more pure population is required (Fig. 4a).

a. Sort ∼0.5 × 10^5^ – 1.0 × 10^5^ cells per well for re-seeding into a 24-well plate.
b. After sorting for CRISPRoff and gRNA expressing cells, the entire population will consist of transfected cells, making it unnecessary to normalize to transfection efficiency at Day 3. This approach results in a cleaner experiment, with nearly all the cells having the gene silenced (Fig. 4b, c), leading to more consistent results compared to using unsorted cells.

#### Considerations and troubleshooting

1. If transfections are low from the CRISPRoff or sgRNA plasmid (<20% BFP+/mCherry+), we recommend optimizing the seeding density and ensuring that the cells are evenly distributed throughout the well. Generally, we have experienced poor transfection efficiencies if the cells are too confluent (error in cell counting) or if cells are localized to the periphery of the well. After seeding the cells in a plate, tilt the plate to ensure that the cells do not attach to the outer edge of the well.
2. If performing plasmid transfection into a different cell type or plate size, we recommend following *Trans*IT-LT1 Transfection Reagent manufacturer’s protocol.
3. As an alternative to transient expression of sgRNA plasmids, we recommend stable expression of the sgRNA vector via lentiviral transduction.
4. Optimization of sgRNA plasmid transfections may be required if using other sgRNA expression vectors.
5. While we provide an example of silencing endogenous genes with a reporter (GFP), antibody staining can be used in cases where no reporter line is available.
6. CRISPROFF mRNA *IN VITRO* SYNTHESIS

Messenger RNA (mRNA) delivery is an effective way to deliver CRISPR-based genome editing tools into mammalian cells (Haasteren et al., 2020; Madigan et al., 2023). Unlike DNA-based delivery systems, mRNA delivery can result in higher transfection efficiency and reduced cellular toxicity (Rohner et al., 2022). Furthermore, there is a reduced risk of insertional mutagenesis with mRNA-based delivery systems, making it an ideal candidate for mRNA therapeutics (Kowalski et al., 2019). Direct delivery of mRNA encoding CRISPRoff into the cytoplasm utilizes the translational machinery of cells for short term, transient expression of CRISPRoff.

**Figure 3.**
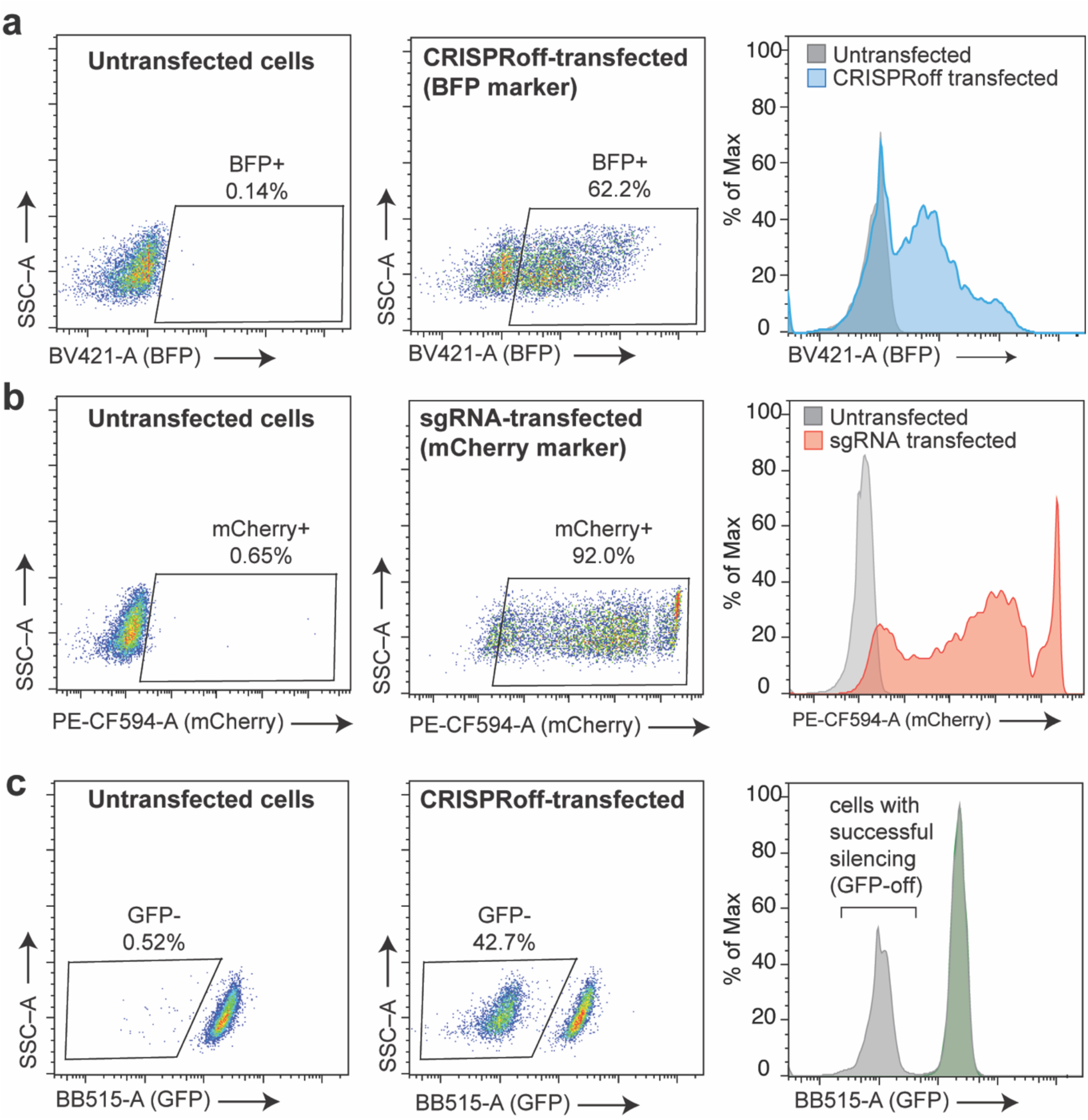
Flow cytometry to measure transfection efficiency and loss of CLTA-GFP as a measure of successful gene silencing. (a) FlowJo software was used to analyze the flow cytometry data two days (Day 3 in protocol) post transfection. (Left) SSC versus BV421-A (BFP) was used to set the gate for CRISPRoff positive cells based on control samples. (Middle) BFP+ gate was used to determine the percent CRISPRoff positive (BFP+) cells from a control sample (cells above diagonal line are BFP+). (Right) Histogram of Left and Middle displaying transfection efficiency of CRISRPoff (BFP). (b) Similar as in (c) but for gating transfection efficiency of gRNA using PE-CF594-A (mCherry). (c) Example flow cytometry data measuring gene knockdown of reporter gene (CLTA) 21 days post transfection using BB515-A (GFP). (Left) SSC versus BB515-A (GFP) was used to set the gate for GFP negative cells based on control samples. (Middle) GFP-gate was used to determine the percent of CLTA negative (GFP-) for unsorted cells (cells below the diagonal line are GFP-). (Right) Histogram of Left and Middle displaying silencing efficiency of CRISPRoff (gray) compared to control sample (green).

**Figure 4.**
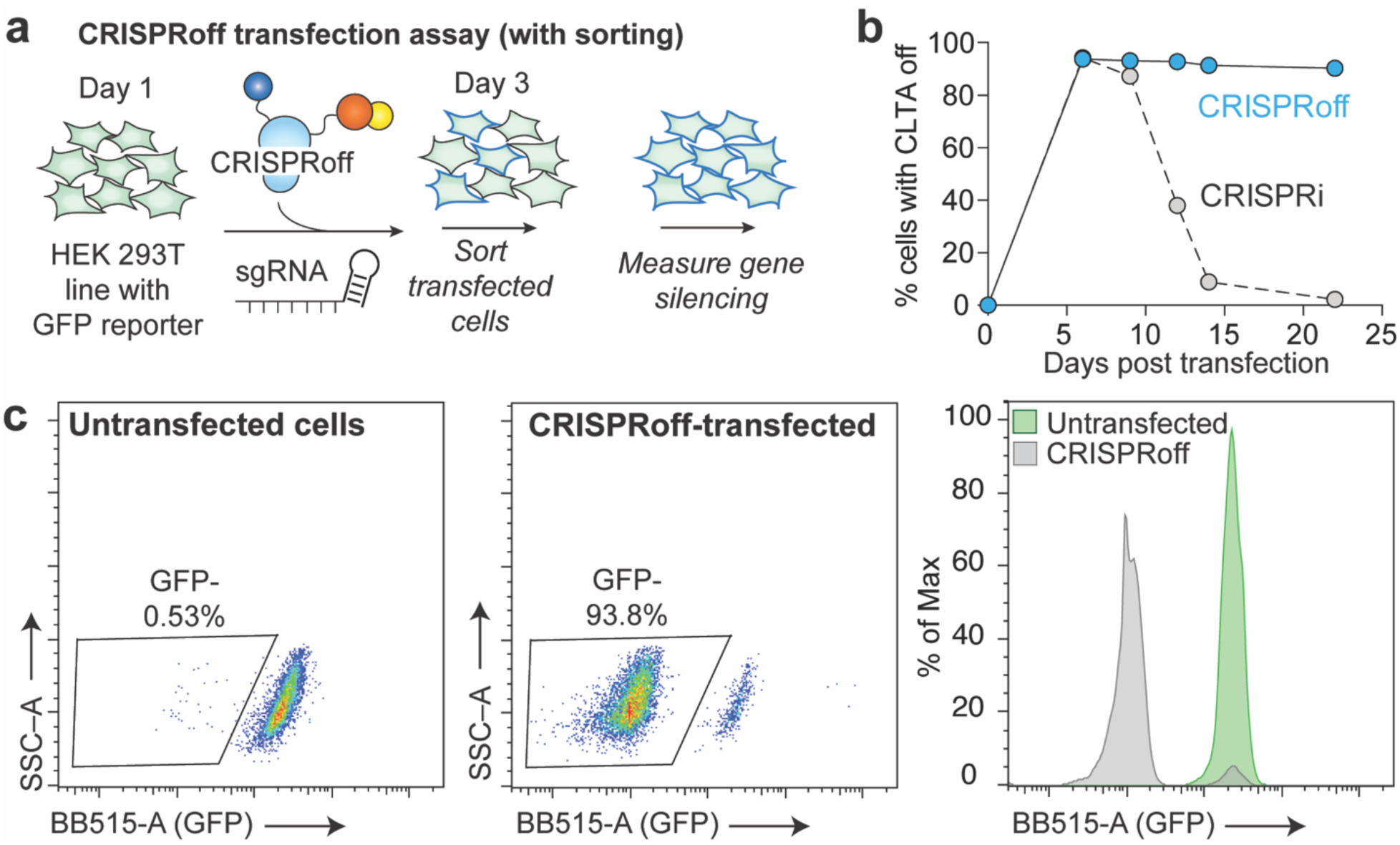
CRISPRoff plasmid transfection assay in HEK 293T cells with sorting for transfected cells (a) Workflow for targeting CRISPRoff to endogenous gene tagged with GFP in HEK 293T cells with sorting transfected cells. (b) A time course analysis of CLTA silencing in HEK 293T cells following CRISPRoff transfection highlighting heritable gene silencing of CRISPRoff (blue) compared to CRISPRi (gray) (c) Similar as in (Fig. 3c) but cells were sorted at two days (Day 3 in protocol) post transfection for CRISPRoff and gRNA expressing cells.

Previously, CRISPRoff has been synthesized as IVT mRNA and delivered into cells to successfully silence genes (Fig. 5) (Nuñez et al., 2021). Here, we provide a further optimized and detailed protocol for synthesizing CRISPRoff mRNA, followed by examples of transfecting and nucleofecting CRISPRoff mRNA into mammalian immortalized cell lines and primary T cells. CRISPRoff mRNA is ∼7,000 nucleotides long, which exceeds the recommended length for many *in vitro* synthesis kits. To optimize both synthesis and delivery, we recommend using the IVT backbone from Aldevron (pALD-CV42 [T7]), the mMessage mMachine™ T7 ULTRA Transcription Kit (Thermo Fisher Scientific) for the IVT reaction and performing enzymatic capping and poly (A) tailing reactions.

**Figure 5.**
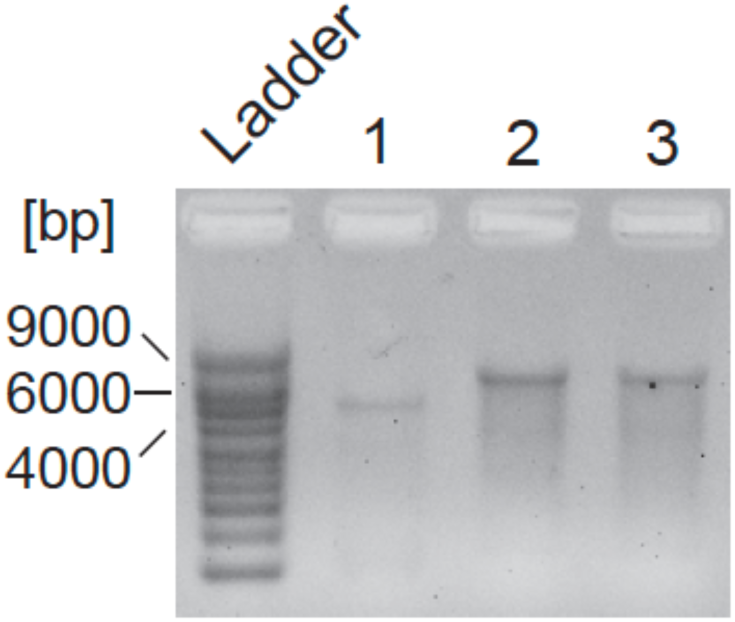
Agarose gel electrophoresis post glyoxal denaturation of IVT mRNA to visualize mRNA size and quality. Lane 1 shows Cas9 mRNA (∼4.2 kb), Lane 2 shows CRISPRoff mRNA (∼7.2 kb), and Lane 3 shows CRISPRoff DNMT3A catalytic mutant mRNA from Nuñez et al., 2021 (∼7.2 kb).

### 5.1. Required materials

- CRISPRoff mRNA vector (generated by cloning CRISPRoff fusion gene (Addgene #167981) into pALD-CV42 [T7] (Aldevron)
- mMessage mMACHINE™ T7 ULTRA Transcription Kit (Thermo Fisher Scientific #AM1345)
- RNaseZap™ RNase Decontamination Solution (Thermo Fisher Scientific #AM9780)
- RNase-free Microfuge Tubes (Thermo Fisher Scientific #AM12400)
- XhoI restriction enzyme (New England Biolabs #R0146S)
- 10X rCutSmart™ Buffer (New England Biolabs #B6004S)
- Thermocycler
- DNA Clean & Concentrator Kit (Zymo Research #D4033)
- Nuclease-free water
- 70% ethanol
- PIPES, free acid
- bis-TRIS [bis-(2-hydroxyethyl) aminotris(hydroxymethyl) methane]
- EDTA
- Nanodrop spectrophotometer
- Qubit™ RNA High Sensitivity (HS) Assay Kit (Invitrogen #Q32852)
- Ambion NorthernMAX glyoxal loading dye Thermo Fisher Scientific (Invitrogen #AM8551)
- Millennial RNA Markers Thermo Fisher Scientific (Invitrogen #AM7150)
- 15 ml conical tubes
- Nuclease free eppendorf tubes
- Pipettes and tips
- Micro-centrifuge

### 5.2. *In vitro* synthesis of CRISPRoff mRNA

#### 5.2.1 Linearization of plasmid

1. Day 1: Assemble the digestion reaction by adding the following to a PCR tube.

2. Incubate the reaction in a thermocycler at 37 °C overnight.

| Component | Amount |
| --- | --- |
| CRISPRoff mRNA vector | 8 ug |
| 10X rCutSmart™ Buffer | 5 µl |
| XhoI restriction enzyme | 2 µl |
| Nuclease-free water | to 50 µl |

#### 5.2.2 Concentration of linearized CRISPRoff DNA template

1. Day 2: Use the DNA Clean and Concentrator Kit to concentrate the DNA. Perform all centrifugation steps at a speed of 10,000-16,000 ×g

a. Divide the linearized CRISPRoff DNA template reaction mixture into two Eppendorf tubes, adding 25 μl of the mixture into each tube.
b. Add 300 μl DNA Binding Buffer to each tube and mix by vortexing.
c. Transfer the mixtures to spin columns placed in collection tubes and centrifuge for 30 s. Discard the flowthrough.
d. Next, add 200 μl of DNA Wash Buffer to each column and centrifuge for 30 s. Repeat this wash step.
e. For elution, use warm (50 °C) nuclease-free water. First, add 6 μl of nuclease-free water directly to the center of each column and incubate at room temperature for 1 min. Then, transfer the column to a 1.5 ml RNase-free Eppendorf tube and centrifuge for 30 s to collect the DNA. Repeat the elution by adding 4 μl of nuclease-free water to the center of the column, incubating for 1 min, and centrifuging for 30 s.
f. Combine the eluates from both columns to obtain a total of 20 μl of linearized DNA. Measure the DNA concentration using a Nanodrop spectrophotometer.

#### 5.2.3 In vitro transcription of CRISPRoff mRNA

1. Spray RNaseZap™ RNase Decontamination Solution on bench, pipettes, tube racks, and gloves to remove RNase contamination.
2. We recommend assembling the transcription reaction with the mMessage mMACHINE™ T7 ULTRA Transcription Kit.
3. Thaw T7 2X NTP/ARCA, 10X T7 Reaction Buffer, and T7 Enzyme Mix on ice.
4. Add the following to a sterile PCR tube. The total reaction volume should equal 20 μl.

5. Gently flick the PCR tube or pipette up and down to mix, then spin the tube briefly to collect reaction mixture at the bottom of the tube.
6. Incubate the transcription reaction at 37 °C for 2 h.
7. Add 1 μl TURBO DNase to the reaction mixture to the transcription reaction to remove trace quantities of DNA contamination. Mix well and incubate the reaction at 37 °C for 15 min.

| **Component** | **Amount** |
| --- | --- |
| Linearized CRISPRoff mRNA vector | 1 µg |
| T7 2X NTP/ARCA | 10 µl |
| 10X T7 Reaction Buffer | 2 µl |
| Nuclease-free water | to 20 µl |

#### 5.2.4 Poly (A) tailing reaction

1. Thaw 5X *E*-PAP Buffer, 25 mM MnCl_2_, and ATP Solution on ice.
2. For the Poly (A) tailing reaction, add the following.

3. Set aside 2.5 μl of the reaction mixture to use as a negative control for the RNA glyoxal gel.
4. Add 4 μl E-PAP to the rest of the above reaction mixture.
5. Gently flick the PCR tube or mix by pipetting up and down, then spin the tube briefly to collect the reaction mixture at the bottom of the tube.
6. Incubate the transcription reaction at 37 °C for 45 min, then place on ice.

| **Component** | **Amount** |
| --- | --- |
| IVT Reaction mixture from above | 20 µl |
| Nuclease-free water | 36 µl |
| 5X E-PAP Buffer | 20 µl |
| 25 mM MnCl<sub>2</sub> | 10 µl |
| ATP Solution | 10 µl |

#### 5.2.5 Lithium chloride precipitation

1. Move the reaction mixture to a 1.5 ml RNase-free eppendorf tube.
2. Add 50 μl of LiCl Precipitation Solution to the reaction mixture and mix thoroughly.
3. Chill the reaction at -20 °C overnight.

#### 5.2.6 CRISPRoff mRNA recovery and quantification

1. Day 3: Centrifuge the reaction mixture at maximum speed at 4 °C for 15 min.
2. Discard the supernatant, taking care not to disturb the mRNA pellet.
3. Wash the mRNA pellet with 1 ml of 70% ethanol and spin at 4 °C for 15 min.
4. Remove ethanol, air dry the mRNA pellet briefly, and resuspend the pellet in 60 μl of nuclease-free water.
5. Measure mRNA concentration by nanodrop spectrophotometer or Qubit™ RNA High Sensitivity (HS) Assay Kit.

#### 5.2.7 CRISPRoff mRNA visualization and quality check

In this protocol, we chose to visualize the *in vitro* synthesized mRNA using denaturing gel electrophoresis (Fig. 5). To ensure the quality and length of the synthesized mRNA, we use glyoxal to denature and visualize CRISPRoff mRNA by size fractionation. About 1 µg of CRISPRoff mRNA is needed to efficiently visualize the mRNA on a gel. This step can be performed in conjunction with quantification of mRNA to ensure that the mRNA is of good quality prior to use. For best results, the synthesized mRNA must be always stored on ice before preparation for gel electrophoresis.

1. Prepare about 500 ml of BPTE buffer for casting and running the mRNA gel. 10X BPTE buffer PIPES, free acid (100 mM) Bis-Tris, free base (300 mM) EDTA (10 mM) [fill up to 500 ml using ddH_2_O] Adjust the pH of the final solution to 6.5 (See Note 1 and 2).
2. Clean the gel casting tray using distilled water.
3. Pour 1% agarose gel in 1X BPTE buffer by adding agarose powder and dissolving it in the microwave (See Note 3, 4).
4. Let the gel set at room temperature to solidify.
5. Prepare the mRNA sample by mixing about 1 μg of CRISPRoff mRNA with glyoxal loading dye (Invitrogen #AM8551) at 1:1 volume.
6. Add 2.5 μl of the saved mRNA reaction mixtures (prior to poly A tail reaction without E-PAP) to glyoxal loading dye at 1:1 volume to serve as a negative control.
7. To prepare the ladder, mix glyoxal loading dye with Invitrogen Millennial RNA Markers (Invitrogen #AM7150) at 1:1 ratio and bring to a desired volume.
8. Incubate the ladder and samples at 50 °C for 30 min, ensuring not to excessively heat the mRNA which can lead to its degradation.
9. Move the samples to ice quickly after incubation to maintain the denatured state of the mRNA (See Note 5).
10. Load the samples in the gel.
11. Run the gel in 1X BPTE buffer at 25 V for 50 min (See Note 6).
12. Image the gel in the ethidium bromide channel to compare RNA length and degradation.
13. Store the synthesized mRNA at -80 °C.

#### Notes

1. It is essential to maintain the pH of the BPTE buffer at acidic pH to prevent reversing of glyoxal reactions.
2. Oakton pH700 pH meter was used to adjust the pH of the solution for accurate measurements.
3. No staining reagents are used in casting the gel.
4. Make sure to use a clean autoclaved flask for casting RNA gels.
5. Denatured mRNA can be stored on ice for 2 – 5 min prior to loading onto the gel.
6. Running the gel too fast can lead to smearing of mRNA, the gel can be run at lower voltages (e.g., 10 V) to prevent smearing.

#### Considerations and troubleshooting

1. Although we recommend the mMessage mMachine™ T7 ULTRA Transcription Kit from Thermo Fisher Scientific for the IVT reaction, other commercially available IVT may also be used, though they may require optimization.
2. We have observed that newly synthesized mRNAs exhibit superior performance compared to those subjected to long-term storage. To mitigate the effects of degradation, it is advisable to avoid frequent freeze-thaw cycles by preparing smaller aliquots of CRISPRoff mRNA.
3. While IVT of CRISPRoff mRNA is generally highly reproducible, consider sourcing high-quality mRNAs from vendors like Aldevron and TriLink® for experiments requiring high consistency, such as in the case with in vivo studies.

## 6. TARGETED GENE SILENCING IN HEK 293T CELLS USING CRISPROFF mRNA

In this section, we describe the nucleofection of IVT synthesized CRISPRoff mRNA into HEK 293T cells that constitutively express an sgRNA targeting the promoter of the endogenous *CLTA* gene, which is tagged with a GFP reporter. The aim of this experiment is to evaluate the efficiency of CRISPRoff mRNA in transcriptionally silencing *CLTA*. By directly introducing CRISPRoff mRNA into the cells, we offer an alternative approach to plasmid DNA-based methods that achieves comparable gene silencing efficiency with potentially reduced cellular toxicity. The primary readout for this experiment is the reduction in CLTA-GFP expression, which can be quantitatively measured in single cells using flow cytometry (Fig. 6a). Additional detection techniques, such as qPCR or Western Blot, may also be employed to verify the extent of CLTA knockdown at the transcript and protein levels respectively.

**Figure 6.**
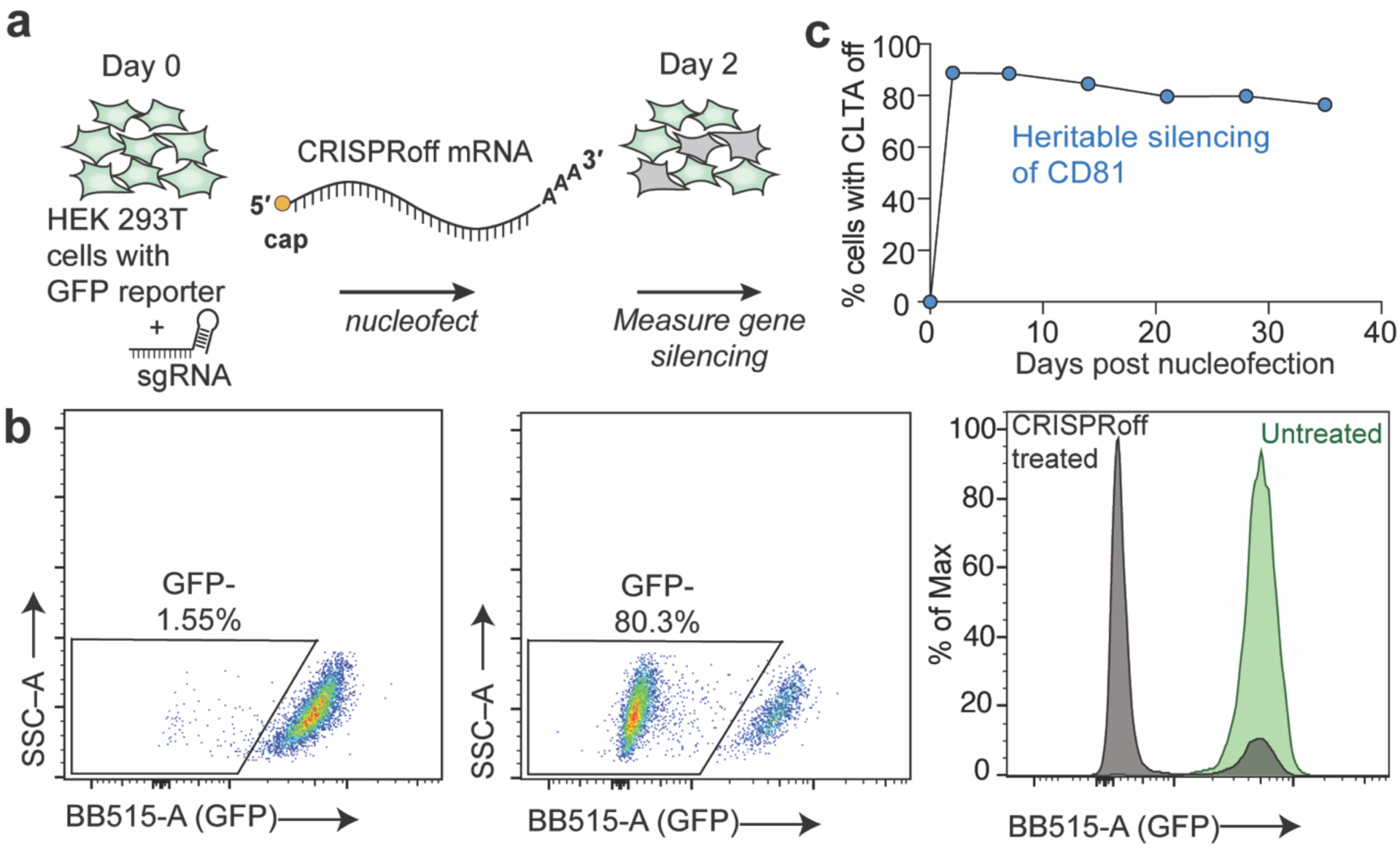
CRISPRoff mRNA nucleofection assay in HEK 293T cells. (a) A schematic illustrating the workflow for nucleofecting HEK 293T cells with CRISPRoff mRNA, targeting the endogenous CLTA gene tagged with GFP. (b) A representative flow cytometry plot showing CLTA silencing (using BB515-A for GFP) in HEK 293T CLTA-GFP cells, 35 days post-nucleofection with CRISPRoff mRNA. (c) A time course analysis of CLTA silencing in HEK 293T cells following CRISPRoff mRNA nucleofection.

### 6.1. Required materials

- HEK 293T cells (ATCC #CRL-3216)
- Dulbecco’s modified eagle medium (DMEM) High Glucose (Thermo Fisher Scientific #11965118)
- 10% FBS (VWR #89510-186)
- Penicillin-Streptomycin-Glutamine (100X) (Gibco #10378016)
- CRISPRoff mRNA (IVT, see Section 5.2)
- SF Cell Line 4D-Nucleofector™ X Kit S (Lonza #V4XC-2032)
- 96-well Shuttle™ Device (Lonza)
- Nucleocuvette™
- Fluorochrome-conjugated antibodies suitable for use in flow cytometry
- BD FACSymphony A1 Cell Analyzer (BD Biosciences)
- DPBS (1X) (Gibco #2027-04)
- Trypan Blue
- Hemocytometer or automated cell counter
- 24-well plates
- CO_2_ incubator
- Centrifuge
- Sterile Eppendorf tubes
- 15 ml conical tubes
- Pipettes and tips

### 6.2. CRISPRoff mRNA nucleofection in HEK 293T cells

Prior to nucleofection, maintain HEK 293T cells in DMEM with 10% FBS and 1X Penicillin-Streptomycin-Glutamine. Passage the cells every 2–3 days, ensuring that they remain at a confluency of 60-70%.

#### 6.2.1 Nucleofection of CRISPRoff mRNA into HEK 293T cells

1. Day 0: Thaw and mix 2 µg of IVT synthesized CRISPRoff mRNA in a sterile RNase-free microcentrifuge tube. Store the mRNA on ice until nucleofection. Ensure that the volume of mRNA does not exceed 10% of the total nucleofection volume (∼22 μl).
2. Warm the SF Cell Line Nucleofector™ solution to room temperature for 15 min before nucleofection. For each reaction, supplement 16.4 μl of SF solution with 3.6 μl of its supplement.
3. Harvest HEK 293T cells and determine the cell count. For each nucleofection reaction, aliquot ∼2.0 × 10^5^ cells into a sterile microcentrifuge tube or use a 15 ml conical tube for multiple reactions.
4. Centrifuge the cells at 500 ×g for 5 min at room temperature and discard the supernatant. Wash the cells once with room temperature PBS at 500 ×g for 5 min, then discard the supernatant.
5. Resuspend the cells in the appropriate volume (18-20 μl) of SF Cell Line Nucleofector™ solution. Add the CRISPRoff mRNA to the resuspended cell solution. For example, mix 2 μl of the mRNA solution with cells resuspended in 18 μl of the SF Cell Line Nucleofector™ solution. Transfer the cells and mRNA mixture into a Lonza Nucleocuvette™, carefully avoiding the formation of bubbles. Gently tap the Nucleocuvette™ to ensure all cells are at the bottom of the Nucleocuvette™.
6. Nucleofect the cells using the Nucleofector™ program CM-130 on the 4D-Nucleofector™ System.
7. Following nucleofection, add 80 μl of the appropriate culture medium to each Nucleocuvette™ well. Transfer ∼100 μl of the cell suspension into a well of a 24-well plate containing 400 μl of pre-warmed culture medium.
8. Incubate the plate at 37 °C with 5% CO_2_ for 24 h.

#### 6.2.2 Assessing gene silencing by flow cytometry

Day 2 and beyond: Unlike plasmid transfections, BFP expression is not detected following nucleofection of CRISPRoff mRNA. Instead, we recommend assessing the efficiency of CRISPRoff gene silencing by directly measuring target protein knockdown using flow cytometry or Western Blot and/or assessing mRNA knockdown using qPCR.

1. Dissociate the nucleofected HEK 293T cells according to the manufacturer’s protocol and measure fluorescence of CLTA-GFP on a flow cytometer.
2. For CRISPRoff mRNA delivery experiments, gene knockdown can be observed as early as day 2 post nucleofection, typically peaking between days 5 and 7 and in most cases, silencing remains heritable and stable over time (Fig. 6b).
3. Furthermore, we recommend additional assessments of target gene silencing 2-3 weeks after nucleofection of CRISPRoff mRNA (Fig. 6c) to assess the durability of long-term gene silencing.

#### Troubleshooting and tips

1. Always include a positive control such as an eGFP/mCherry reporter mRNA to assess nucleofection efficiency.
2. Include a non-targeting sgRNA control to assess specificity of gene silencing.
3. We recommend testing 3 – 5 guides per gene target and considering the pooling of multiple guides to enhance efficiency.
4. In this example, the sgRNA is stably expressed in the HEK 293T cell line. If a stable HEK 293T cell line expressing the target sgRNA is unavailable, consider co-nucleofecting the sgRNA along with CRISPRoff mRNA. Based on our experience, co-nucleofection of CRISPRoff mRNA (2 μg per nucleofection reaction) and sgRNA (2-3 μg per nucleofection reaction) per 2.0 × 10^5^ cells also results in robust gene silencing.
5. While we provide an example of silencing endogenous genes with a reporter (GFP), antibody staining can be used in cases where no reporter line is available.
6. If nucleofection-related toxicity is observed, consider increasing the number of cells per nucleofection reaction.
7. In some cases, optimizing CRISPRoff mRNA (2 – 4 µg per nucleofection reaction) per 2.0 × 10^5^ cells may be necessary to achieve the desired effective dose.
8. If gene silencing leads to cellular toxicity, consider seeding the cells in a 96-well plate instead of a 24-well plate post-nucleofection.
9. Although our protocol is optimized for the SF Cell Line 4D-Nucleofector™ X Kit S, further optimization of cell number and CRISPRoff mRNA amounts will be necessary when scaling up to more cells.
10. For cell types that are less suitable for nucleofection, we suggest encapsulating CRISPRoff mRNAs in commercial or in vitro-formulated lipid nanoparticles (LNPs) to enhance transfection efficiency and improve delivery.

## 7. TARGETED GENE SILENCING IN hTERT RPE-1 CELLS USING CRISPROFF mRNA

In this section, we outline the transfection of IVT synthesized CRISPRoff mRNA into hTERT RPE-1 cells, which stably express an sgRNA targeting the promoter of the endogenous *CD81* gene. hTERT RPE-1 cells, derived from human retinal pigment epithelial cells, maintain a near-diploid karyotype, making them a more physiologically relevant model for studying normal cellular processes (Hindul et al., 2022). The objective of this experiment is to assess the effectiveness of CRISPRoff mRNA in transcriptionally silencing *CD81*. The primary readout for this experiment is the reduction in CD81 expression, which can be quantitatively assessed in single cells through CD81 protein staining and flow cytometry (Fig. 7a). Additional methods, such as qPCR or Western Blot, may also be used to confirm the extent of CD81 knockdown at the transcript and protein levels respectively.

**Figure 7.**
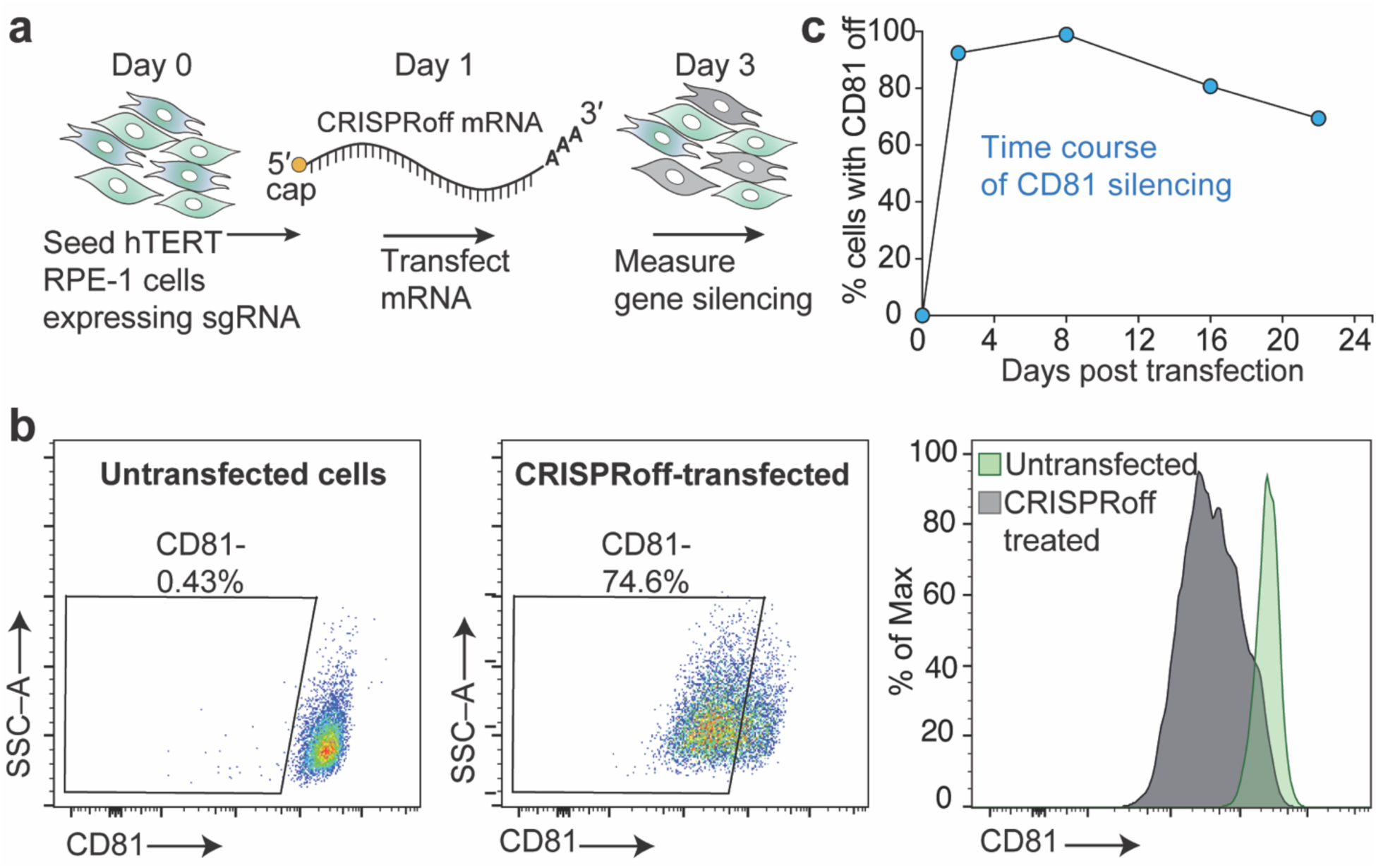
CRISPRoff mRNA transfection assay in hTERT RPE-1 cells. (a) A schematic illustrating the workflow for transfecting hTERT RPE-1 cells with CRISPRoff mRNA. (b) A representative flow cytometry plot showing CD81 silencing in hTERT RPE-1 cells, 2 days post-transfection with CRISPRoff mRNA. (c) A time course analysis of CD81 silencing in hTERT RPE-1 cells following CRISPRoff mRNA transfection.

### 7.1. Required materials

- hTERT RPE-1 cells (ATCC #CRL-4000)
- DMEM/F-12, GlutaMAX™ supplement (Thermo Fisher Scientific #105650-018)
- 10% FBS (VWR #89510-186)
- Penicillin-Streptomycin-Glutamine (100X) (Gibco #10378016)
- CRISPRoff mRNA (IVT, see Section 5.2)
- CRISPRoff *CD81* sgRNA as detailed in Section 3.2 (Ordered from IDT or Synthego)
- Lipofectamine™ MessengerMax™ Transfection Reagent (Invitrogen #LMRNA008)
- Fluorochrome-conjugated antibodies suitable for use in flow cytometry
- BD FACSymphony A1 Cell Analyzer (BD Biosciences)
- Trypan Blue
- Hemocytometer or automated cell counter
- 24-well plates
- CO_2_ incubator
- Centrifuge
- Sterile Eppendorf tubes
- 15 ml conical tubes
- Pipettes and tips

### 7.2. CRISPRoff mRNA transfection in hTERT RPE-1 cells

Prior to the transfection, maintain human hTERT RPE-1 cells in DMEM/F-12 with 10% FBS and 1X Penicillin-Streptomycin-Glutamine. Passage the cells every 2–3 days, ensuring that they remain at a confluency of 60-70%.

#### 7.2.1 Seeding hTERT RPE-1 cells for CRISPRoff mRNA transfection

1. Day 0: Seed 1 × 10^5^ hTERT RPE-1 cells in a 24-well plate containing 0.5 ml of the appropriate cell culture medium. Adjusting the seeding density might be necessary depending on the timing of transfection and the method used for cell counting.

#### 7.2.2 Transfection of CRISPRoff mRNA into hTERT RPE-1 cells

1. Day 1: Under a microscope, check seeded cell density to see if cells are ∼60 – 70% confluent.
2. Bring Opti-MEMTM and MessengerMAX™ Reagent to room temperature ∼30 min before transfection.
3. In an RNase-free sterile microcentrifuge tube, for each transfection reaction, dilute 1.5 μl of MessengerMAX™ Reagent in 25 μl of pre-warmed Opti-MEM™ Medium. Mix thoroughly and incubate the transfection reaction at room temperature for 10 min.
4. Mix 1 μg of CRISPRoff mRNA with 25 μl of pre-warmed Opti-MEM™ Medium in a sterile microcentrifuge tube.
5. Combine the diluted mRNA mixture with the diluted MessengerMAX™ Reagent, mix well, and incubate at room temperature for 5 min.
6. Add ∼50 μl of mRNA-lipid mixture dropwise to the cells.
7. Incubate the plates at 37 °C with 5% CO2 for 24 h.

#### 7.2.3 Assessing gene silencing by flow cytometry

Day 3 and beyond: Unlike plasmid transfections, BFP expression is not detected following transfection of CRISPRoff mRNA. Instead, we recommend assessing the efficiency of CRISPRoff gene silencing by directly measuring target protein knockdown using flow cytometry or Western Blot and/or assessing at the RNA level using qPCR.

1. Dissociate the transfected hTERT RPE-1 cells according to the manufacturer’s protocol, perform antibody staining for CD81, and then measure CD81 knockdown using a flow cytometer.
2. For CRISPRoff mRNA delivery experiments, gene knockdown can be observed as early as day 3 (Fig. 7b), typically peaking between days 6 and 8 and in most cases, silencing remains heritable and stable over time.
3. Furthermore, we recommend additional assessments of target gene silencing 2-3 weeks after nucleofection of CRISPRoff mRNA (Fig. 7c) to assess the durability of long-term gene silencing.

#### Troubleshooting and tips

1. Always include a positive control such as eGFP or mCherry reporter mRNA to evaluate transfection efficiency.
2. Include a non-targeting sgRNA control to assess specificity of gene silencing.
3. We recommend testing 3 – 5 guides per gene and considering the pooling of multiple guides to enhance the efficiency of CRISPRoff-mediated silencing.
4. Consider increasing cell numbers per transfection reaction to account for nucleic acids mediated toxicity.
5. In some cases, optimization of CRISPRoff mRNA (1 – 3 μg per reaction) may be necessary to achieve the desired effective dose.
6. When scaling reactions to different volumes than those that we recommend, it is important to optimize the volumes of transfection reagents and the amounts of CRISPRoff mRNA.

## 8. TARGETED GENE SILENCING IN PRIMARY HUMAN DONOR DERIVED T CELLS USING CRISPROFF mRNA

In this section, we describe the co-nucleofection of IVT synthesized CRISPRoff mRNA into primary human T cells, along with an sgRNA targeting the promoter of the endogenous *CD81* gene. Unlike immortalized cell lines, which often harbor genetic abnormalities, primary T cells offer a more accurate and relevant model for studying native immune functions. Derived directly from human peripheral blood mononuclear cells (PBMCs), primary T cells provide a physiologically relevant system, making them ideal for developing effective cancer immunotherapies (Stadtmauer et al., 2020).

The aim of this experiment is to evaluate the effectiveness of CRISPRoff mRNA in transcriptionally and heritably silencing *CD81* in activated primary T cells. We first provide an overview of the process for isolating primary human T cells from donor derived PBMCs and activating them to induce proliferation. In this example, we also outline the steps for co-delivering CRISPRoff mRNA and an sgRNA into primary T cells. This approach is particularly useful because creating stable sgRNA-expressing lentiviral lines in primary cells, such as T cells, can be challenging. The primary readout for this experiment is the reduction in CD81 protein, which can be quantitatively measured by antibody staining for CD81 followed by flow cytometry analysis (Fig. 8a). Additionally, qPCR or Western blot may be employed to verify CD81 knockdown at the transcript and protein levels respectively.

**Figure 8.**
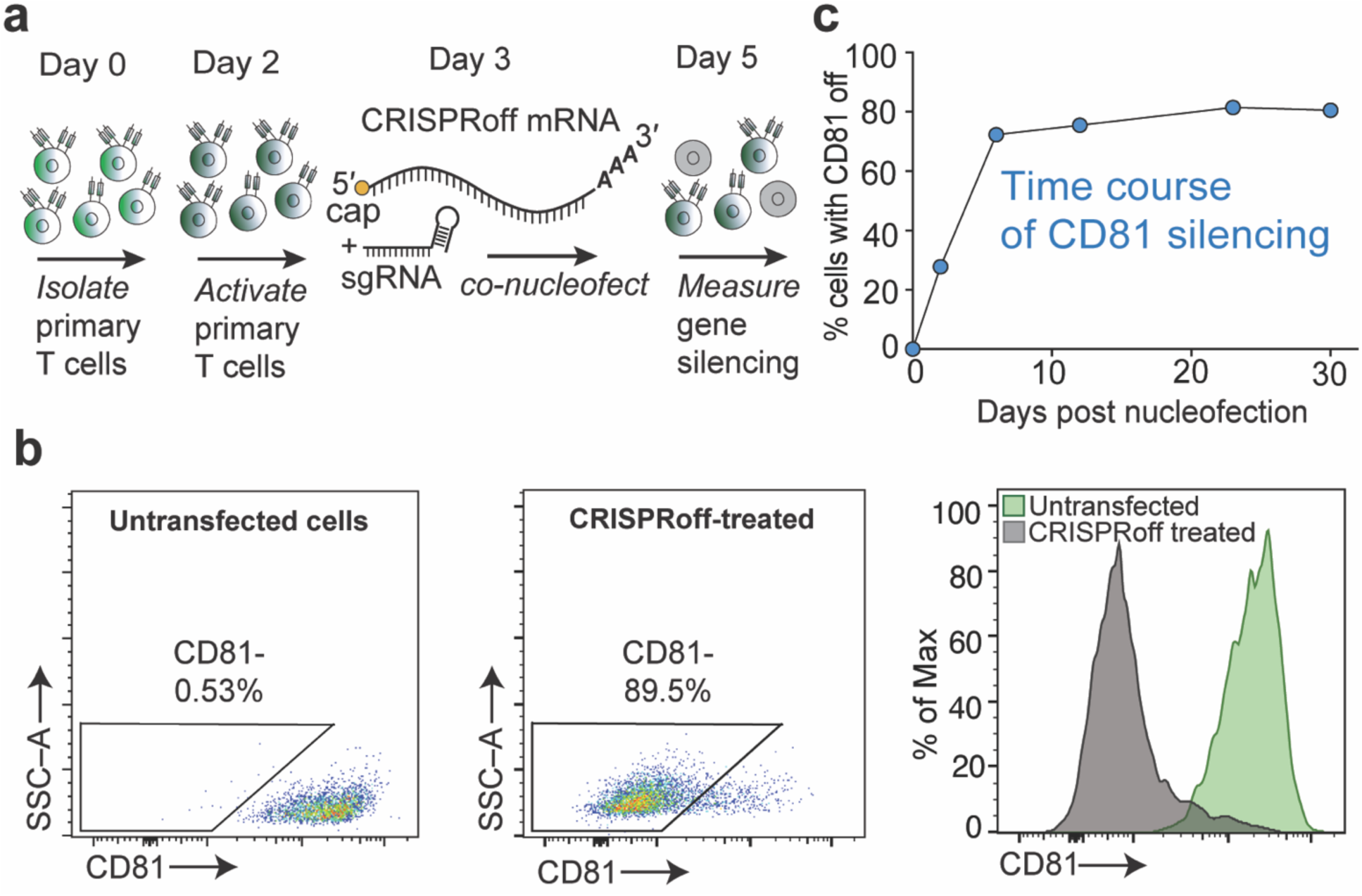
CRISPRoff mRNA co-nucleofection assay in activated primary human T cells. (a) A schematic illustrating the workflow for isolating, activating, and co-nucleofecting primary human T cells with CRISPRoff mRNA and sgRNA. (b) A representative flow cytometry plot demonstrating CD81 silencing in primary human T cells, 16 days post-nucleofection with CRISPRoff mRNA. (c) A time course analysis of CD81 silencing in primary human T cells following CRISPRoff mRNA nucleofection.

### 8.1. Required materials

- Human Peripheral Blood Mononuclear Cells, Frozen (STEMCELL Technologies #70025.3)
- X-VIVO™ 15 Serum-free Hematopoietic Cell Medium (Lonza #02-053Q)
- N-acetyl L-cysteine (Sigma-Aldrich #A7250)
- 2-mercaptoethanol (Gibco #21985023)
- 10% FBS (VWR #89510-186)
- Penicillin-Streptomycin-Glutamine (100X) (Gibco #10378016)
- IL-2 (Peprotech #200-02)
- IL-7 (Peprotech #200-07)
- IL-15 (R&D Systems #BT-015-010)
- EasySep™ magnetic Cell Isolation kit (STEMCELL Technologies #17951)
- Dynabeads™ Human T-Activator CD3/CD28 for T Cell Expansion and Activation (Thermo Fisher Scientific #11131D)
- CRISPRoff mRNA (IVT, see Section 5.2)
- CRISPRoff *CD81* sgRNA as detailed in Section 3.2 (Ordered from IDT or Synthego)
- P3 Cell Line 4D-Nucleofector™ X Kit S (32 RCT) (Lonza #V4XP-3032)
- 96-well Shuttle™ Device (Lonza)
- Nucleocuvette™
- APC anti-human CD81 (TAPA-1) Antibody (Biolegend #349510)
- BD FACSymphony A1 Cell Analyzer (BD Biosciences)
- DPBS (1X) (Gibco #2027-04)
- Trypan Blue
- Hemocytometer or automated cell counter
- 24-well plates
- CO_2_ incubator
- Centrifuge
- Sterile Eppendorf tubes
- 15 ml conical tubes
- Pipettes and tips

### 8.2. CRISPRoff mRNA and sgRNAs co-nucleofection in primary human T cells

#### 8.2.1 Isolation of primary human T cells from PBMCs

1. Day 0: Isolate bulk CD3+ T lymphocytes from healthy donor PBMCs using the EasySep™ Magnetic Cell Isolation Kit, following the manufacturer’s instructions.

#### 8.2.2 Activation of primary human T cells

1. Day 2: Activate primary T cells using Dynabeads™ Human T-Activator CD3/CD28 for T Cell Expansion and Activation as per the manufacturer’s protocol.
2. Culture activated Primary T cells in X-VIVO™ medium containing fetal bovine serum (5%), 2-mercaptoethanol (50 μM), N-acetyl L-cysteine (10 mM), IL-2 (300 U/ml), IL-7 (5 ng/ml), and IL-15 (5 ng/ml).
3. We recommend using freshly isolated and activated primary T cells for CRISPRoff mRNA nucleofection. For cells already in culture, maintain activated primary human T cells by passaging them every 2–3 days and maintaining a cell density of 1 × 10⁶ cells per ml.

#### 8.2.3 Co-nucleofection of CRISPRoff mRNA and sgRNA into activated primary human T cells

1. Day 3: To prepare the CRISPRoff mRNAs-sgRNA mixture, mix 2.5 μg of the target gene sgRNA (*CD81* in this example) and 1 μg of CRISPRoff mRNA in a sterile RNase-free microcentrifuge tube and store on ice until ready for nucleofection. It is recommended that the mRNA-sgRNA mixture should not exceed 10% of the total nucleofection reaction (∼22 μl).
2. Warm up the P3 Primary Cell 4D-Nucleofector™ Solution 15 min prior to nucleofection. For each reaction, mix 16.4 μl P3 Primary Cell 4D-Nucleofector™ Solution with 3.6 μl of its supplement.
3. Harvest activated primary T cells and determine their cell density. For each nucleofection reaction, aliquot 1 × 10^6^ cells into a sterile microcentrifuge tube.
4. Centrifuge the cells at 500 ×g for 5 min at room temperature, then discard the supernatant. Wash the cells once with room temperature PBS at 500 ×g for 5 min and discard the supernatant.
5. Resuspend the cells in the appropriate volume (∼18-20 μl) of P3 Primary Cell 4D-Nucleofector™ Solution. Add the sgRNA-mRNA mixture into the cells. For example, mix 2 μl of sgRNA-mRNA solution with cells resuspended in 18 μl of P3 Primary Cell 4D-Nucleofector™ Solution. Transfer the cells and mRNA-sgRNA mixture into one well of a Lonza Nucleocuvette™, being careful to avoid creating bubbles. Gently tap the Nucleocuvette™ to ensure all the cells settle at the bottom of the plate.
6. Nucleofect the cells using the Nucleofector™ program DS-137 on the 4D-Nucleofector™ System.
7. After nucleofection, add 80 μl of pre-warmed T cell culture medium into each Nucleocuvette™ well. Transfer ∼100 μl of the cell suspension cells into a well of a 96-well plate containing 100 μl of pre-warmed T cell culture medium with growth factors.
8. Incubate the 96-well plate at 37 °C with 5% CO_2_ for 24 h.
9. Replace T cell media and growth factors as needed, approximately every 3-4 days.

#### 8.2.4 Assessing gene silencing by flow cytometry

Day 5 and beyond: Unlike plasmid transfections, BFP expression is not detected following nucleofection of CRISPRoff mRNA. Instead, we recommend assessing the efficiency of CRISPRoff gene silencing by directly measuring target protein knockdown using flow cytometry or Western Blot and/or assessing mRNA knockdown using qPCR.

1. Harvest nucleofected primary human T cells, perform antibody staining for CD81, and then measure CD81 knockdown using a flow cytometer.
2. For CRISPRoff mRNA delivery experiments, gene knockdown can be observed as early as day 5, typically peaking between days 8 and 10 and in most cases, silencing remains heritable and stable over time (Fig. 8b).
3. Furthermore, we recommend additional assessments of target gene silencing 2-3 weeks after nucleofection of CRISPRoff mRNA (Fig. 8c) to assess the durability of long-term gene silencing.

#### Troubleshooting and tips

1. Always include a control eGFP/mCherry reporter mRNA control to assess nucleofection efficiency.
2. Include a non-targeting sgRNA control to determine the specificity of gene silencing.
3. We recommend testing 3-5 guides per gene and considering the pooling of multiple guides to enhance efficiency.
4. For primary cells such as T cells, we highly recommend using commercially available sgRNAs from vendors such as IDT and Synthego.
5. Consider increasing cell numbers per nucleofection reaction to account for nucleofection mediated toxicity.
6. Optimize sgRNA amounts (1-5 µg per reaction) and CRISPRoff mRNA (1-3 µg per reaction) per 1 million cells as needed to achieve the desired effective dose.
7. We recommend using freshly isolated T cells for silencing experiments; however, we have also observed stable silencing in primary T cells that were cryopreserved without prior stimulation, then thawed and used in CRISPRoff experiments.
8. Although our protocol is optimized for the P3 Cell Line 4D-Nucleofector™ X Kit S, further optimization of cell number and mRNA amounts will be necessary when scaling up.
9. For cell types that are less suitable for nucleofection, we suggest encapsulating CRISPRoff mRNAs in commercial or in vitro-formulated lipid nanoparticles (LNPs) to enhance transfection efficiency and improve delivery.

## 9. CONCLUSION

CRISPRoff is an emerging tool for heritable gene repression in the rapidly advancing field of epigenetic editing, offering a robust and reversible approach to gene silencing by altering the epigenome. By combining the precise DNA binding ability of CRISPR systems with the power to edit epigenetic marks, CRISPRoff serves as a highly programmable platform for targeted gene perturbation without altering the underlying DNA sequence or inducing DNA breaks. In this protocol, we present a detailed guide for utilizing CRISPRoff to program heritable gene silencing in human cells. Beyond CRISPRoff, other epigenetic editing tools have been reported and our protocol can be applied to the delivery of these technologies into mammalian cells (Alerasool et al., 2022; Cano-Rodriguez et al., 2016; Chavez et al., 2015; DelRosso et al., 2023; Hilton et al., 2015; Kearns et al., 2015; Mahata et al., 2023; McCutcheon et al., 2024; O’Geen et al., 2017; O’Geen et al., 2019; Policarpi et al., 2024; Tanenbaum et al., 2014). These emerging technologies extend the capabilities of CRISPR-based epigenetic editors, allowing researchers to activate or repress genes in a more controlled and distinct manner.

While CRISPRoff can silence a vast majority of genes in the human genome (Nuñez et al., 2021), not all genes are equally amenable to CRISPRoff mediated silencing. The effectiveness of this tool is affected by several factors, and it is important to recognize that CRISPRoff is still in the early stages of development. One key factor to consider is the local epigenetic landscape of the target gene (Yeo et al., 2018), as pre-existing epigenetic marks can potentially hinder or neutralize the effectiveness of CRISPRoff. Additionally, the specific cell type being targeted plays a major role, as genes in different cell types may have distinct epigenetic profiles. For example, primary cells and immortalized cell lines can respond variably to CRISPRoff induced epigenetic editing. Another key variable is the CpG density within the target gene promoter region, as promoter regions with low CpG content may be less susceptible to methylation-based silencing by CRISPRoff. All of these factors must be evaluated for optimal efficacy and success of CRISPRoff mediated gene silencing.

With these considerations in mind, we have shown in this protocol that CRISPRoff shows remarkable versatility across various immortalized cell types, such as HEK 293T and hTERT RPE-1 cells. Moreover, in this paper, we also highlight the efficacy of CRISPRoff in targeting an endogenous gene in primary donor-derived human T cells, thereby expanding its potential for developing potent immunotherapies. The ability of CRISPRoff to induce stable and heritable gene silencing highlights its promising application for *in vivo* epigenetic editing and offers exciting possibilities for its use as a therapeutic platform.

## ACKNOWLEDGMENTS

We thank members of the Nuñez lab for establishing and optimizing the protocols described in the manuscript. We thank Stephen Floor for advice on mRNA *in vitro* synthesis and gel analysis. This work was funded in part by research grants awarded to J.K.N.: a Hanna Gray Fellowship from the Howard Hughes Medical Institute (GT15341), NIH R35 MIRA (R35GM155044-01), CRISPR Cures for Cancer Initiative (Gladstone Institutes, UCB 20240097), a Spark Award from Bakar Labs (51474), and an Innovation Award from the Laboratory for Genomics Research (UCB 052611). J.K.N. is an Investigator of the Chan Zuckerberg Biohub San Francisco. R.K.P. is funded by graduate fellowships from the University of California Cancer Research Coordinating Committee and the Shurl and Kay Curci Foundation. I.J.O. is funded by graduate fellowships from the University of California, Berkeley Mentored Research Award, a Gilliam Fellowship from the Howard Hughes Medical Institute, and a National Science Foundation Graduate Research Fellowship. N.S.D. is funded by a postdoctoral fellowship from the California Institute for Regenerative Medicine (CIRM). C.D.N. acknowledges support from the SURF Rose Hills Fellowship from the University of California, Berkeley and the Amgen Scholars Program at the University of California, San Francisco.

